# Decidual immune response following COVID-19 during pregnancy varies by timing of maternal SARS-CoV-2 infection

**DOI:** 10.1101/2021.11.20.469369

**Authors:** Lillian J. Juttukonda, Elisha M. Wachman, Jeffery Boateng, Mayuri Jain, Yoel Benarroch, Elizabeth S. Taglauer

## Abstract

While COVID-19 infection during pregnancy is common, fetal transmission is rare, suggesting that intrauterine mechanisms form an effective blockade against SARS-CoV-2. Key among these is the decidual immune environment of the placenta. We hypothesized that decidual leukocytes are altered by maternal SARS-CoV-2 infection in pregnancy and that this decidual immune resonse is shaped by the timing of infection during gestation. To address this hypothesis, we collected decidua basalis tissues at delivery from women with symptomatic COVID-19 during second (2^nd^ Tri COVID, n=8) or third trimester (3^rd^ Tri COVID, n=8) and SARS-CoV-2-negative controls (Control, n=8). Decidual natural killer (NK) cells, macrophages and T cells were evaluated using quantitative microscopy, and pro- and anti-inflammatory cytokine mRNA expression was evaluated using quantitative reverse transcriptase PCR (qRT-PCR). When compared with the Control group, decidual tissues from 3^rd^ Tri COVID exhibited significantly increased macrophages, NK cells and T cells, whereas 2^nd^ Tri COVID only had significantly increased T cells. In evaluating decidual cytokine expression, we noted that IL-6, IL-8, IL-10 and TNF-α were significantly correlated with macrophage cell abundance. However, in 2^nd^ Tri COVID tissues, there was significant downregulation of IL-6, IL-8, IL-10, and TNF-α. Taken together, these results suggest innate and adaptive immune responses are present at the maternal-fetal interface in maternal SARS-CoV-2 infections late in pregnancy, and that infections earlier in pregnancy show evidence of a resolving immune response. Further studies are warranted to characterize the full scope of intrauterine immune responses in pregnancies affected by maternal COVID-19.

## 1. Introduction

Healthy pregnancies have been challenged by the pandemic of COVID-19, the illness caused by SARS-CoV-2. Disease severity correlates with expression of proinflammatory cytokines, sometimes characterized as a ‘cytokine storm’(Song *et al.*, 2020). Pregnant women contract COVID-19 at rates similar to the general population and develop a spectrum of illness ranging from asymptomatic carriage to critical illness (Dubey *et al.*, 2020; Allotey *et al.*, 2020), but extremely severe COVID-19 and death are more likely in pregnant women than nonpregnant women (Zambrano *et al.*, 2020). Infection during pregnancy is also associated with an increased risk for preterm birth, preeclampsia, stillbirth, and low birth weight (Wei *et al.*, 2021).

However, vertical transmission of SARS-CoV-2 from mother to fetus during pregnancy is rare (Tolu, Ezeh and Feyissa, 2021; Juan *et al.*, 2020; Kotlyar *et al.*, 2021; Raschetti *et al.*, 2020; Al-Matary *et al.*, 2021; Facchetti *et al.*, 2020) with positive neonatal nasopharyngeal PCR test rates ranging between 0.5 and 2.5%. Multiple studies have now substantiated that placental tissues express the SARS-CoV-2 receptor ACE-2 (Taglauer *et al.*, 2020; Li *et al.*, 2020; Pavličev *et al.*, 2017; Edlow *et al.*, 2020; Ouyang *et al.*, 2021). SARS-CoV-2 has also been detected in placental tissues (Taglauer *et al.*, 2020; Sharps *et al.*, 2020; Kotlyar *et al.*, 2021) and is able to infect placental trophoblast cells *in vitro* (Ouyang et al., 2021; Lu-Culligan et al., 2021). Given these findings in conjunction with the persistent rarity of fetal transmission, additional factors within the maternal-fetal interface likely work together with the villous placenta to ultimately prevent the transplacental passage of SARS-CoV-2. Of central importance are intrauterine immune responses governed primarily by maternal leukocytes in the decidual tissues adjacent to the villous placenta. Among these, decidual T cells, NK cells, and macrophages are known to respond to viral infection at the maternal-fetal interface, either through direct cellular or indirect cytokine-mediated pathogen specific responses (Parker *et al.*, 2020; Jabrane-Ferrat, 2019; Granja *et al.*, 2021; Yockey and Iwasaki, 2018).

Previous studies have identified global inflammatory responses in the placenta (Lu-Culligan *et al.*, 2021) and sexually dimorphic alteration in interferon response genes in villous placental tissues from pregnant individuals with COVID-19 in the third trimester of pregnancy (Bordt *et al.*, 2021). However, the leukocyte and cytokine profiles of the decidual-specific immune response to maternal SARS-CoV-2 require more specific characterization, particularly when infection occurs prior to the third trimester.

We hypothesized that maternal immune cells in the decidua respond to COVID-19 via changes in immune cell number and production of cytokines and that these changes can be impacted by the timing of maternal infection in pregnancy. To test this hypothesis, we evaluated decidual leukocytes and cytokine profiles of decidual tissues from women with symtomatic COVID-19 infections in their 2^nd^ or 3^rd^ trimesters of pregnancy as compared to controls to characterize the trajectory of the decidual immune responses against SARS-CoV-2 at the maternal-fetal interface.

## 2. Materials and Methods

### 2.1 Study enrollment

#### Setting

Boston Medical Center (BMC), the largest urban safety-net hospital in New England. All patient enrollment and tissue collection was approved by the Boston University Medical Campus Institutional Review Board.

#### Retrospective cohort

An initial retrospective cohort (April-May 2020) involved tissue collection from placentas designated for pathology analysis either from women who tested positive via SARS-CoV-2 nasopharyngeal PCR testing at the time of delivery or from contemporary controls who tested negative. For this cohort, informed consent was waived, and samples were accompanied by limited demographic data available from chart review.

#### Prospective cohort

Mother-infant dyads were consented in a prospective cohort study (July 2020-April 2021). Inclusion criteria included age minimum of 18 years, singleton pregnancy, and English/Spanish speaking. For the COVID-19 group, mothers had a positive nasopharyngeal PCR test for SARS-CoV-2 infection at any point during pregnancy. For the control group, mothers must not have had a documented SARS-CoV-2 infection at any point during pregnancy.

#### Data collection

Demographic and clinical variables (as summarized in Tables 1 and 2) were obtained from the electronic medical record (EMR) and recorded in a secure, de-identified RedCap Database (https://www.project-redcap.org).

**Table 1:**
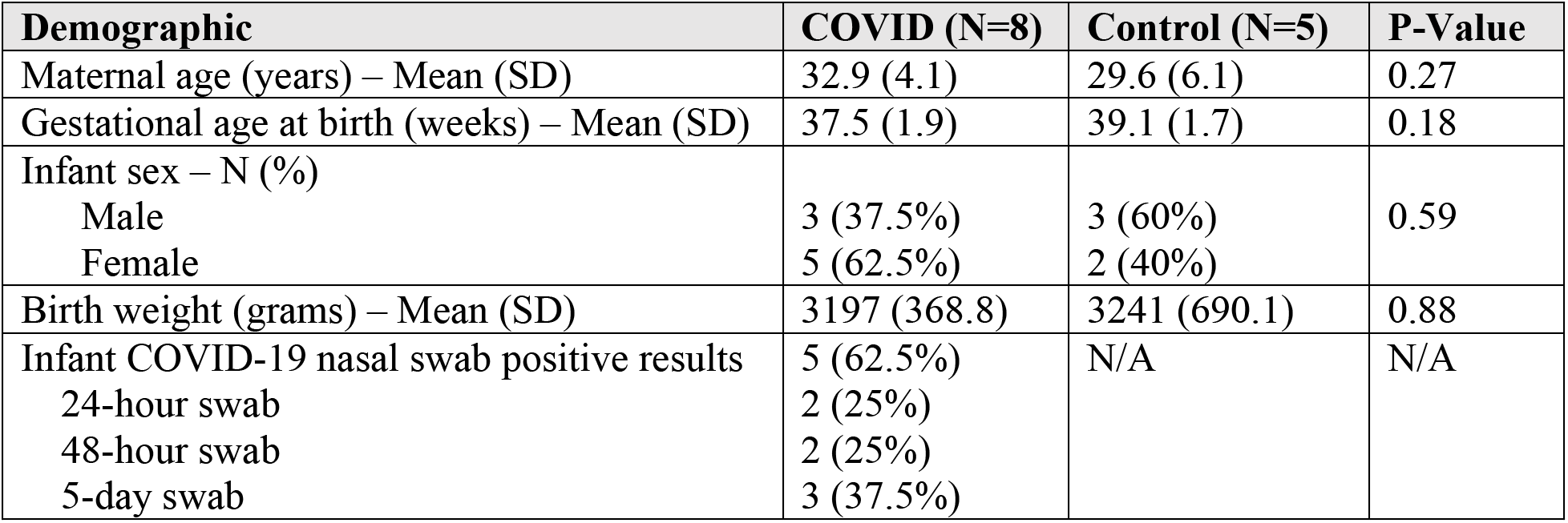
Demographic data from retrospective COVID-19 cohort.

| **Demographic** | **COVID (N=8)** | **Control (N=5)** | **P-Value** |
| --- | --- | --- | --- |
| Maternal age (years) – Mean (SD) | 32.9 (4.1) | 29.6 (6.1) | 0.27 |
| Gestational age at birth (weeks) – Mean (SD) | 37.5 (1.9) | 39.1 (1.7) | 0.18 |
| Infant sex – N (%) |  |  |  |
| Male | 3 (37.5%) | 3 (60%) | 0.59 |
| Female | 5 (62.5%) | 2 (40%) |  |
| Birth weight (grams) – Mean (SD) | 3197 (368.8) | 3241 (690.1) | 0.88 |
| Infant COVID-19 nasal swab positive results | 5 (62.5%) | N/A | N/A |
| 24-hour swab | 2 (25%) |  |  |
| 48-hour swab | 2 (25%) |  |  |
| 5-day swab | 3 (37.5%) |  |  |

**Table 2:**
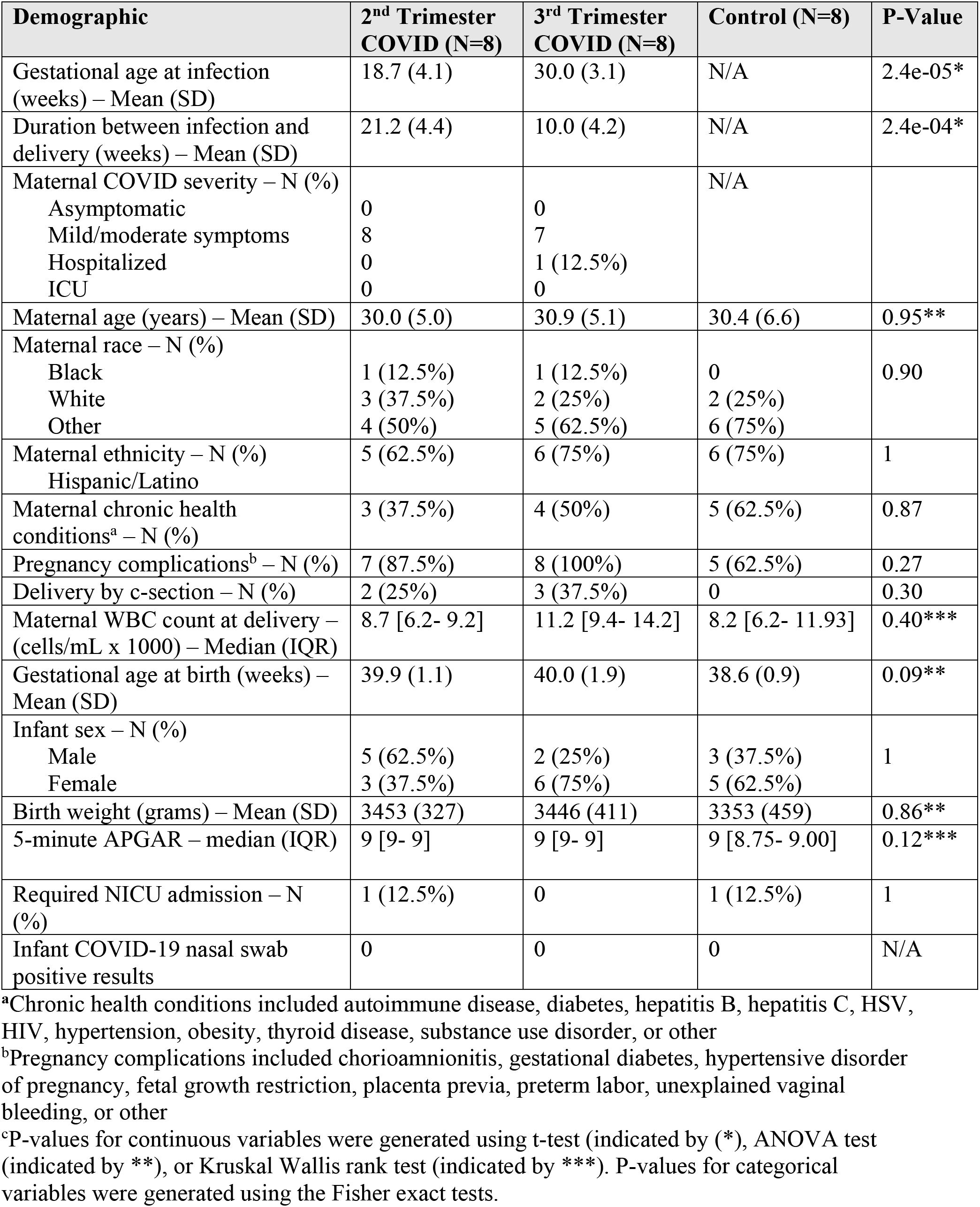
Demographic and clinical data from prospective COVID-19 cohort.

### 2.2 Sample collection

#### Retrospective cohort

Tissues were placed in 4% buffered formalin at time of delivery and evaluated by board-certified pathologists. Full thickness placental biopsies were dissected from the remaining formalin fixed placental tissues, soaked in 18% sucrose, embedded in optimal cutting temperature (OCT) compound (Fisher) and frozen at −80°C. Ten micron cryosections were subjected to antigen retrieval (Sodium citrate, 0.05% Tween 20) prior to immunostaining.

#### Prospective cohort

Fresh placental tissue was collected and processed within 6 hours after delivery. Dissected decidual tissue biopsies were flash-frozen with dry ice and stored at −80°C. Placental full-thickness biopsies were dissected and fixed with formaldehyde/sucrose for 72 hours, embedded in OCT (Fisher) and frozen at −80°C. Tissues were cryosectioned at 10 μM thickness.

### 2.3 Immune Cell Analysis

#### Immunohistochemistry

Placental tissue sections were washed with PBS-T, permeabilized, and incubated with Histostain Plus Broad Spectrum Blocking Solution (#859043, Novex by LifeTechnologies). Primary antibodies were diluted 1:100 in PBS-T as follows: *single labeling*: CD14 (mouse anti-human, 14-149-82, Invitrogen), CD56 (mouse anti-human, 14-0567-82, Invitrogen); *double labeling*: SARS CoV-2 spike glycoprotein (rabbit anti-human, ab272504 Abcam), CD3 (mouse anti-human, 14-0038-82, Invitrogen). Fluorescently labeled secondary antibodies were diluted 1:500 in PBS-T as follows: *single labeling:* Alexa 594 (anti-mouse, ab150108 Abcam); *double labeling*: AlexaFluor 594 (anti-mouse, ab150108 Abcam), AlexaFluor 647 (anti-rabbit, A21244, LifeTechnologies). Control slides were incubated with secondary antibodies alone. Washed slides were cover slipped with Prolong Gold with DAPI (ThermoFisher) and cured overnight prior to imaging. All cohort slides were stained in bulk and imaged within 24-48 hours of staining.

#### Microscopy

Images were acquired on a Nikon deconvolution wide-field epifluorescence microscope using NIS-Elements Software (Nikon). Decidua basalis areas were manually surveyed at 100× followed by automated acquisition at 200× of 4 tiled images from 5 randomized areas per slide. Exposure times were standardized for each target and remained constant throughout image acquisition to ensure accurate comparative quantification.

#### Quantitative Image Analysis

Image area and integrated density were measured via ImageJ software (imagej.net) for each immunofluorescent 200× image (n=5/slide) along with mean fluorescence values from 5 randomly selected background readings per image cohort were used to calculate a corrected total cell fluorescence (CTCF) per published protocols (Taglauer *et al.*, 2020; Benarroch *et al.*, 2021; Kenan *et al.*, 2020). An average CTCF was calculated from secondary only negative control slides (n=3 per staining assay) and used to calculate a final Fluorescence Ratio: target antigen CTCF/secondary only control CTCF. All ImageJ analysis and calculations were performed on blinded samples.

### 2.4 Gene Expression Analysis

#### RNA Isolation and cDNA synthesis

RNA was isolated from frozen placental decidual biopsies (50-75 μg) using the RNAqueous™ kit (#AM1912, Invitrogen). RNA quantity and purity was assessed using a Nanodrop™ spectrophotometer (ThermoFisher). cDNA was generated from equal amounts of RNA (800 ng) using a RevertAid First Strand cDNA synthesis kit (ThermoFisher).

#### qRT-PCR

PrimePCR assays for IL-1β, IL-6, IL-8, IL-10, IFN-γ, TNF-α, and GAPDH (#10025636, BIO-RAD) and iTaq™ Universal SYBR Green Supermix (#1725120, BIO-RAD) utilizing a BIO-RAD thermocycler with the manufacturer’s recommended settings. Assays were run in technical triplicate and no-reverse transcriptase controls were included to ensure there was no DNA contamination. GAPDH was used to normalize sample abundance. Fold changes were calculated for COVID-19 groups relative to the control group using the standardized ΔΔCT method (Livak and Schmittgen, 2001).

### 2.5 Statistical analysis

#### Clinical and demographic data

Categorical variables were reported as percentages [n (%)] and continuous variables were reported as either mean with standard deviation [mean(sd)] or median with interquartile range [median (IQR)], where appropriate. For Table 1, P-values were generated using t-tests. For Table 2, P-values for continuous variables were generated using t-test (indicated by (*), ANOVA test (indicated by **) or Kruskal Wallis rank test (indicated by ***). P-values for categorical variables were generated using the Fisher exact tests. Differences were considered significant at α = 0.05. Analyses were conducted using R version 3.6.1 (https://www.r-project.org).

#### Laboratory data

P-values for quantitative immunofluorescence were generated using ANOVA with Tukey’s multiple comparison’s test. P-values for fold-changes by qRT-PCR results were generated using the one-sample Wilcoxon Signed Rank Test compared to a theoretical median of 1.0. Pearson test was used to generate P-values for correlations between cytokine fold-changes and immune cell abundance. Differences were considered significant at α = 0.05. All laboratory statistical analysis was performed using Prism 9 software (GraphPad).

## 3. Results

### 3.1 Decidual macrophages and NK cells are increased in 3^rd^ Trimester COVID 19 infections

We first assessed decidual leukocytes in the retrospective cohort. There were no significant differences in gestational age at birth, birth weight, or maternal age between the COVID-19 infected (n=8) and control (n=5) groups (**Table 1**). Of note, multiple infants in this cohort tested positive for SARS-CoV-2 by nasopharyngeal PCR after birth (**Table 1**). In comparison to controls, decidua basalis samples from women with COVID-19 had significantly increased macrophage (**Figure 1A and B**) and NK cell populations (**Figure 1C and D**). There were no differences between female and male infants (data not shown).

**Figure 1:**
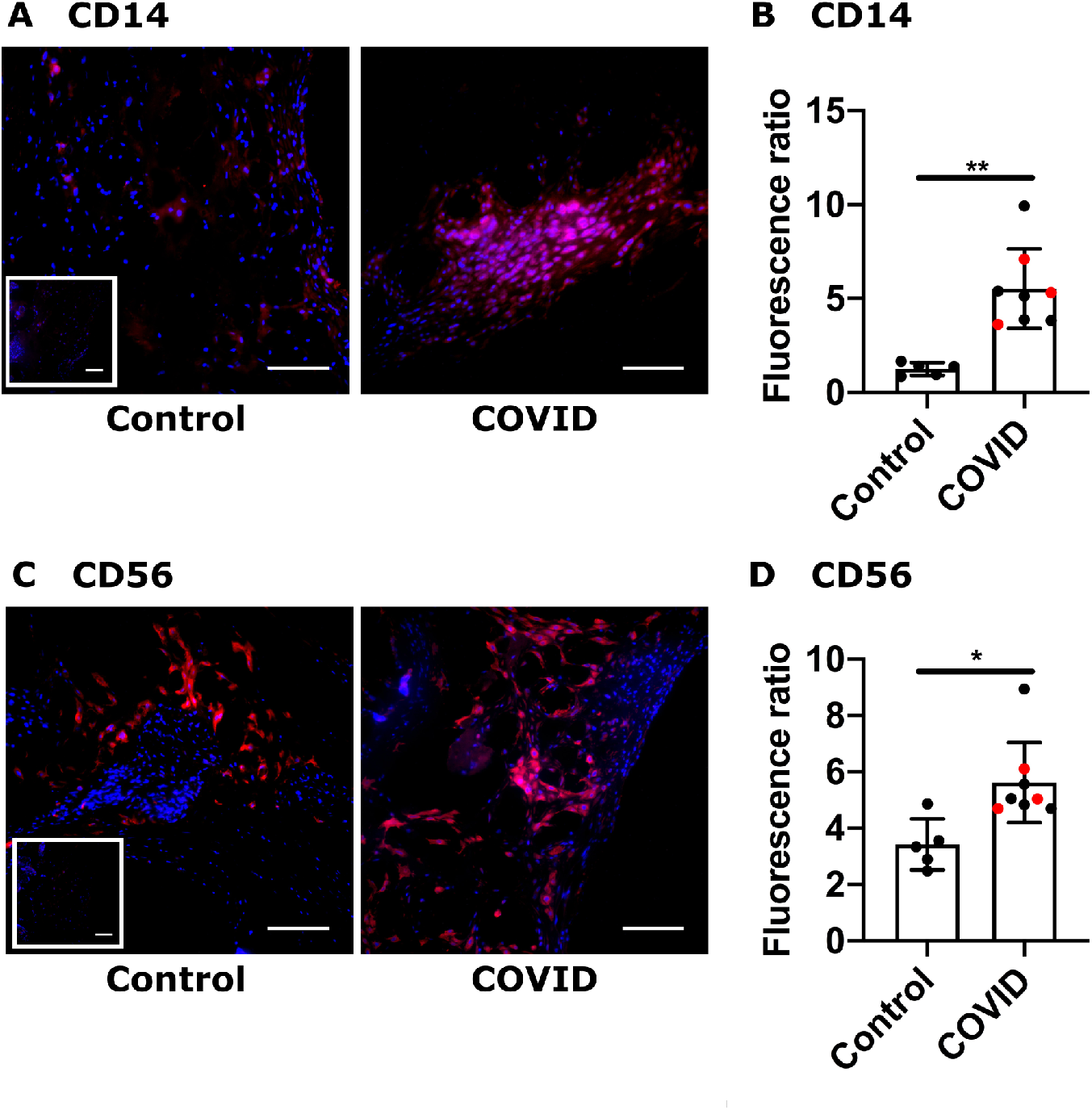
Placental maternal immune cell infiltrates from retrospective COVID-19 cohort. **(A, C)** Representative images (200x) of decidual areas stained for **(A)** CD14 (red) or **(C)** CD56 (red) immunofluorescence. **(B, D)** Graphical analysis of comparative fluorescence quantitation of **(B)** CD14 and **(D)** CD56. Red circles indicate dyads with an infant who tested positive for SARS-CoV-2 by nasal PCR swab at 24 or 48 hours of life. SARS-CoV-2 positive (COVID, n = 8) or negative (control, n = 5). * *p* <0.05; ** *p* <0.01. Scale bar: 50μm; insets: secondary-only controls.

### 3.2 The abundance of decidual macrophages, NK cells, and T cells in the differs based on gestational timing of COVID-19 infection

Decidual leukocyte abundance was examined further in our prospective cohort, with 16 total in the COVID-19 and 8 in the control group. All participants with COVID-19 were symptomatic in this study, but only one required hospitalization. There were no significant differences in maternal or infant demographics (**Table 2**). None of the infants in the COVID-19 groups tested positive for SARS-CoV-2 by nasopharyngeal PCR, but notably testing was only performed in mothers with a symptomatic infection at time of delivery. Further, there were no significant differences in placental pathology reports observed in the COVID-19 compared to control groups (**Supplemental Table 1**).

We next assessed whether there was evidence of SARS-CoV-2 infection in the placenta at time of delivery by performing immunofluorescence for the Spike protein. Two placentas from the 3^rd^ Tri COVID and one from the 2^nd^ Tri COVID group stained positive for Spike protein (**Figure 2A**). Evaluation of decidual CD14 staining showed that decidual macrophages were significantly increased in the 3^rd^ Tri COVID group (**Figure 2B & C**), confirming our retrospective study results. In contrast, there was no difference in macrophage cell staining between Control and 2^nd^ Tri COVID groups (**Figure 2B & C**). Analysis of CD56 staining demonstrated a significant increase in decidual NK cells in the 3^rd^ Tri COVID group, whereas there were no differences in NK cell abundance between Control and 2^nd^ Tri COVID groups (**Figure 2 D & E**).

**Figure 2:**
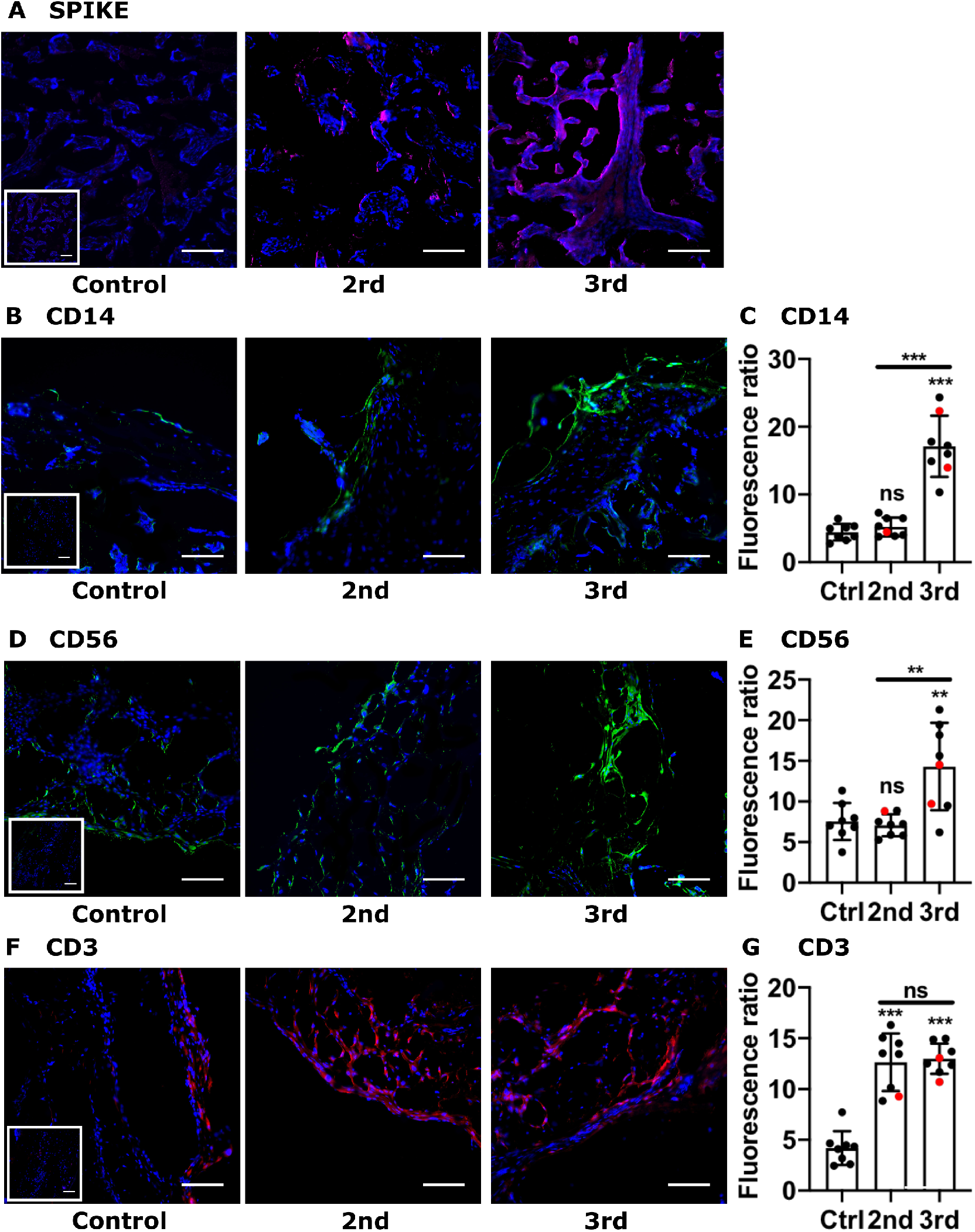
Immune cell infiltrates from prospective COVID-19 cohort. **(A)** Representative immunofluorescence images (200x) of the three placental decidua tissues that stained positive for SARS-CoV-2 SPIKE (red). **(B, D, F)** Representative images (200x) of decidual areas stained for **(B)** CD14 (green), **(D)** CD56 (green), or **(F)** CD3 (red) immunofluorescence. **(C, E, G)** Graphical analysis of comparative fluorescence quantitation of **(C)** CD14, **(E)** CD56, or **(G)** CD3. Red circles indicate placentas with positive staining for SARS-CoV-2 Spike protein. 2^nd^ = 2^nd^ Tri COVID (n = 8); 3^rd^ = 3^rd^ Tri COVID (n = 8); control/ctrl = negative for COVID (n = 8). ** *p* <0.01; ** *p* < 0.001. Scale bar: 50μm; insets: secondary-only controls.

Similar to macrophage and NK cells, CD3^+^ T cell staining was significantly increased in the 3^rd^ Tri COVID group (**Figure 2F & G**). However, in contrast to macrophage and NK cell staining, the T cell infiltrate was also increased in 2^nd^ Tri COVID as compared to Control group (**Figure 2F & G**). Finally, in evaluating gender specific changes, there was a small, but significant, difference in CD14 staining comparing male and female infants in Control but not COVID-19 groups (**Supplemental Figure 1**). NK cells and T cells did not show any significant differences in abundance based on gender (Data not shown).

### 3.3 Decidual cytokine expression is positively correlated with macrophage and NK cell abundance

We next chose to evaluate a targeted panel of cytokines which had been identified in SARS-CoV-2 immune responses in other organ systems (Tang *et al.*, 2020; Leisman *et al.*, 2020) and cytokines known to be to be present within the maternal-fetal interface (Mori *et al.*, 2016). For cytokine evaluation, we first conducted a correlation analysis between cytokine expression (fold change relative to Control tissues) and decidual leukocyte abundance in the COVID groups. We found that there was a significant positive correlation between IL-6, IL-8, IL-10, and TNF-α expression and macrophage abundance, and a significant positive correlation between TNF-α and NK cell abundance (**Figure 3)**. No significant correlations were found among cytokine expression and T cell abundance (**Supplemental Figure 2**).

**Figure 3:**
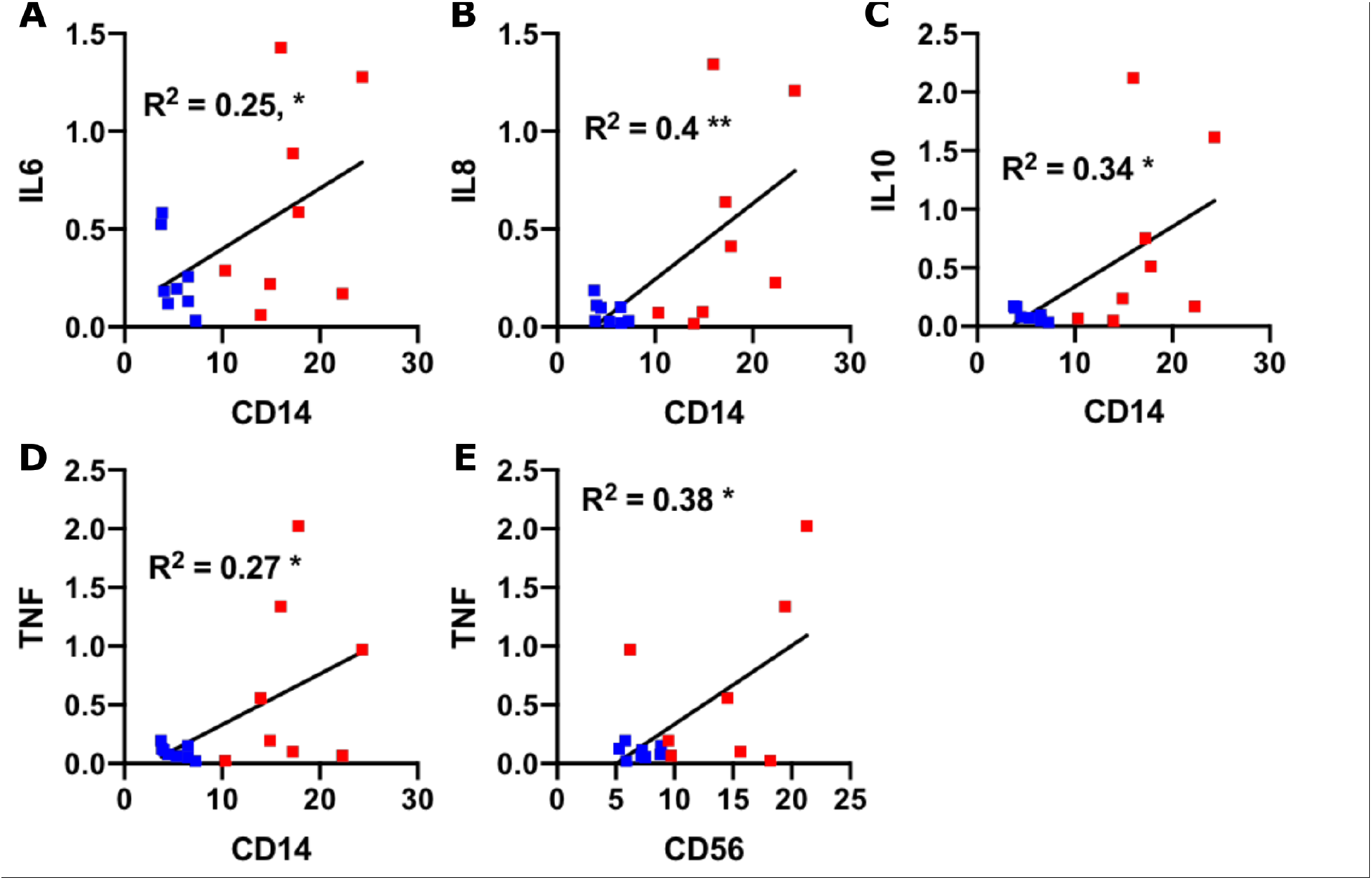
Correlation between cytokine abundance and immune cells. **(A-E)** Scatter plots of the indicated cytokines (Y axis; fold change relative to control) and immune cell markers (X axis; fluorescence ratio) for each placental sample (COVID-19+; n=16). Blue symbols = 2^nd^ Tri COVID; red symbols = 3^rd^ Tri COVID. *, *p* < 0.05; **, *p* < 0.01.

### 3.4 IL-6, IL-8, IL-10 and TNF-α are downregulated in decidual tissues from 2^nd^ trimester COVID-19 infections

We next compared total cycokine changes in 2^nd^ Tri COVID and 3^rd^ Tri COVID groups (evaluated as fold change relative to Control tissues). Interestingly, in the 2^nd^-Tri COVID group, there was a strong pattern of transcript downregulation, with significant decrease in IL-6, IL-8, IL-10, and TNF-α and no change in abundance for IL-1β or IFN-γ (**Figure 4**). Further, in the 3^rd^-Tri COVID tissues, there were no differences in expression of IL-1β, IL-6, IL-10, TNF-α, or IFN-γ and a significant downregulation of IL-8 (**Supplemental Figure 3**). Finally, there were no differences in cytokine levels based on the gender of the infant (data not shown).

**Figure 4:**
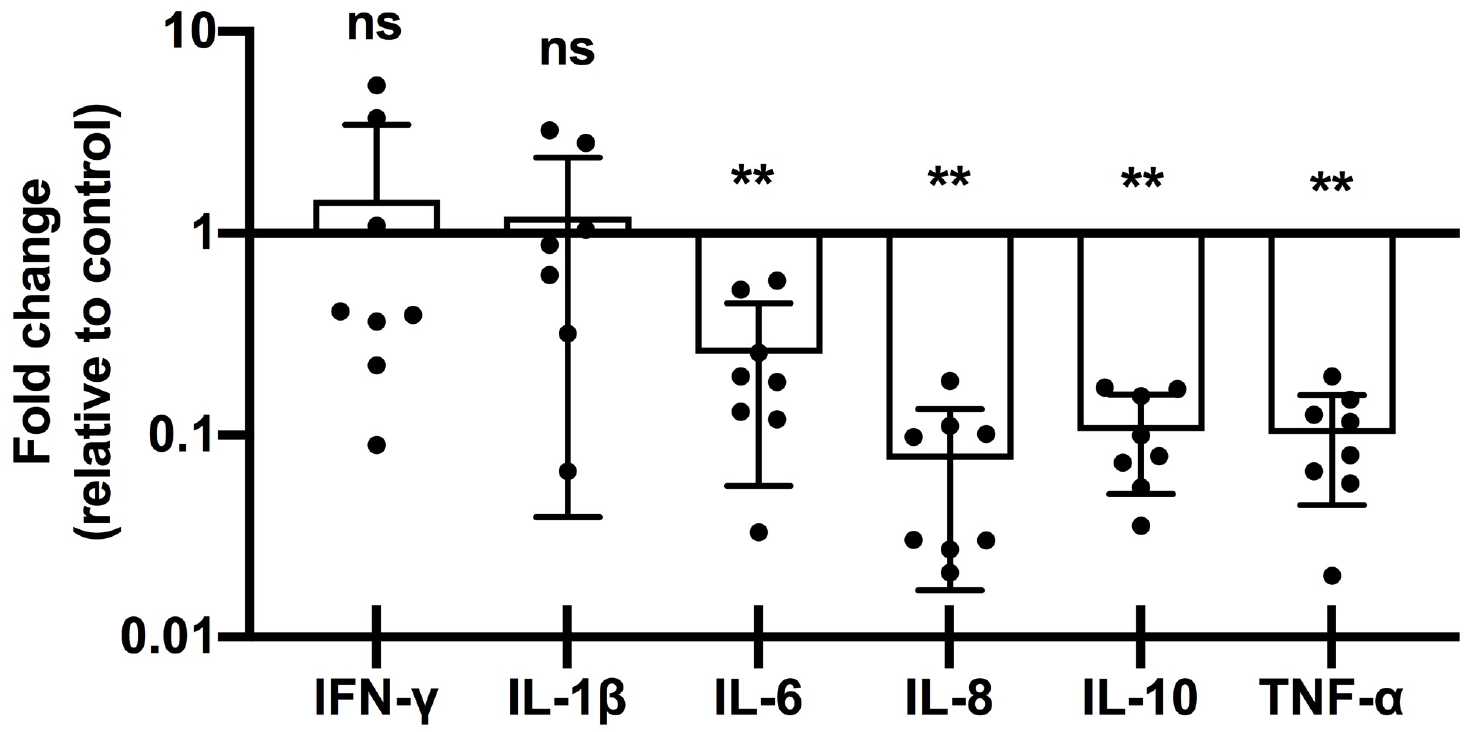
Placental cytokine mRNA downregulation in 2^nd^ Tri COVID group. Fold-changes in cytokine abundance for IFN-γ, IL-1β, IL-6, IL-8, IL-10, and TNF-α by qRT-PCR of placental decidua from women with 2^nd^ trimester COVID infection (2nd, n = 8) calculated as fold-change compared to control (n = 8). ns = not significant; ** *p* <0.01.

## 4. Discussion

In the current study, we identified an association between COVID-19 in pregnancy and an increased immune cell abundance in the decidua basalis, with the specific cell types dependent on gestational timing of SARS-CoV-2 infection. Further, we identified that levels of multiple cytokine transcripts correlated with macrophage abundance, which was driven by 3^rd^ trimester COVID-19 infections. In contrast, there was an overall trend of downregulation in the pro and anti-inflammatory cytokine production in pregnancies with 2^nd^ trimester maternal COVID-19 disease. Taken together, these results provide new insignt into the the immune responses at the maternal-fetal interface and highlight the importance of evaluating various gestational stages of maternal COVID-19 infections to fully characterize the scope of the intrauterine immune response against SARS-CoV-2.

Our findings of inflammatory cells in the decidua is consistent with our previously reported findings in this same cohort identifying gross pathology of intervillous and subchorionic fibrosis (Taglauer *et al.*, 2020). There is an association between COVID-19 in the 3^rd^ trimester and increased macrophage, NK cell, and T cell populations, whereas COVID-19 in the 2^nd^ trimester was associated only with increased T cell populations. Importantly, this study demonstrates an association between COVID-19 infection and increased immune cells that is dependent on the gestational age at which infection occurred. The inclusion of a second-trimester infection group is particularly valuable, as there is very limited knowledge of second-trimester SARS-CoV-2 infection and long-term immune consequences at the maternal-fetal interface. The finding that increased T cell infiltrates were present both 10 weeks (3^rd^ Tri COVID group) and 21 weeks (2^nd^ Tri COVID group) following infection is intriguing. Further studies to phenotypically characterize the T cells by subtype would help to reveal the function of these cells. Effector cells may restrict active viral replication, whereas memory cells may protect the fetus from repeat attacks and regulatory cells may promote resolution of inflammation.

In our cytokine analysis, we anticipated that pro-inflammatory cytokines would be elevated in the decidual tissues from women with third-trimester COVID-19, but instead we found no significant differences with the exception of IL-8 downregulation. While these results differ from an early report of global placental inflammatory signaling in placental tissues from pregnancies with 3^rd^ trimester COVID-19 infections (Lu-Culligan *et al.*, 2021), our findings are similar to more recently published data that there were no changes associated with COVID-19 in pregnancy at the placental level for TNF-α, IL-1β, or IL-6 and upregulation of IFN-γ was only seen in male fetuses (Bordt *et al.*, 2021). It is possible that our study missed the window of pro-inflammatory cytokine upregulation, as infections in the third trimester group were on average 10 weeks prior to delivery. An alternative hypothesis is that the decidual immune response to COVID-19 does not involve upregulation of these pro-inflammatory cytokines, which may be beneficial to the developing fetus as there are pathologies associated with increased levels of IFN-γ, IL-1β, IL-6, and TNF-α in mouse models (Yockey and Iwasaki, 2018).

In tissues where cytokine abundance increased compared with control tissues, there was a positive correlation primarily with macrophage abundance and IL-6, IL-8, IL-10, and TNF-α expression. Macrophages are known to produce these cytokines in response to stimulation via the Toll-like-receptors and other pathogen-associated molecular pattern receptors (Hoo, Nakimuli and Vento-Tormo, 2020); it is plausible that this association is due to production of these cytokines by maternal macrophages. We also identified a positive correlation with TNF-a and NK cell abundance, which is also consistent with the known role of decidual NK cells in viral responses in pregnancy (Jabrane-Ferrat, 2019).

Interestingly, proinflammatory cytokines were downregulated in the placenta following second-trimester COVID-19. This may be because the tissues were in the resolution phase post infection, but other explanations include that the T-cell population observed in the second trimester decidua is exhibiting an exhaustion phenotype with decreased production of cytokines. Future studies should interrogate these cells for T cells markers of exhaustion, particularly PD-1 as this is known to be a prominent signaling pathway in decidual T cells at the maternal-fetal interface (Taglauer *et al.*, 2008).

The current study has several limitations. First, we only had access to basic clinical demographic data from the retrospective cohort, and it is possible that the women had severe COVID-19 disease as these tissues were collected during the first-wave of COVID-19, limiting the generalizability of these results. Importantly, the findings of increased macrophages and NK cells were replicated in our third-trimester group in the prospective, fully-consented cohort study with contemporary controls. A limitation in the prospective cohort study is the small sample size collected from a single clinical site.

Taken as a whole, these results demonstrate that immune cell infiltrates in the placenta following COVID-19 are driven by timing of infection during gestation. While additional studies are required to more fully evaluate the immune profile in pregnancies affected by maternal COVID-19, these data provide early evidence supporting the role of decidual leukocytes in the physiologic response against SARS-CoV-2 at the maternal-fetal interface.

## 5. Acknowledgements

We thank staff of the Boston Medical Center Labor and Delivery Unit, Neonatal ICU, and Newborn Nursery for their help in obtaining clinical samples. Funding: this work was supported by a Boston University Clinical & Translational Science Institute Pilot Grant to E.M.W., a Lovejoy Resident Research Grant to L.J.J., and an American Academy of Pediatrics Resident Research Award to L.J.J.

## 7. Tables

## 8. Figures

**Supplememental Table 1:** Pathology diagnoses from prospective COVID-19 cohort.

| **PATHOLOGY DIAGNOSIS** | **COVID (N=16)** | **CONTROL (N=8)** | **P-Value** |
| --- | --- | --- | --- |
| Meconium Histiocytosis | 7 (58.3%) | 4 (66.7%) | 1.00 |
| Infarcts | 3 (25%) | 2 (33.3%) | 1.00 |
| Chorioamnionitis | 1 (8.3%) | 0 | ---- |
| Fibrin deposition | 3 (25%) | 1 (16.7%) | 1.00 |
| Fetal chorionic vessel vasculitis | 1 (8.3%) | 0 | ---- |
| Hypoplastic villi | 1 (8.3%) | 0 | ---- |
| Acute subchorionitis | 3 (25%) | 2 (33.3%) | 1.00 |
| Decidual vasculopathy | 1 (8.3%) | 0 | ---- |

**Supplemental Figure 1:**
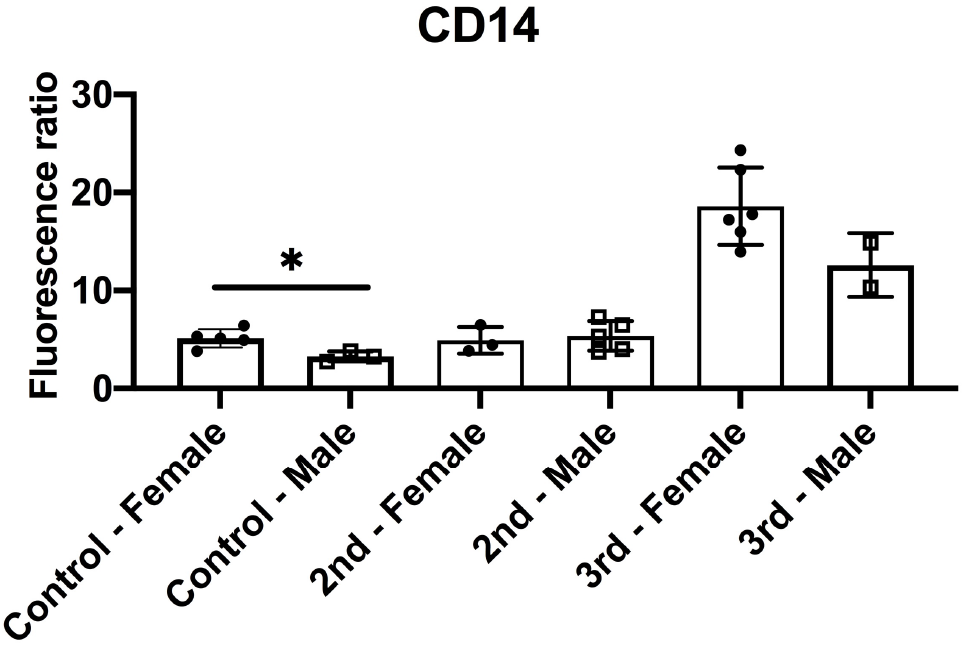
Infant gender sub-analysis of CD14 fluorescence staining in prospective cohort. Data from Figure 2 were divided by infant gender assigned at birth. p < 0.05 by t-test.

**Supplemental Figure 2:**
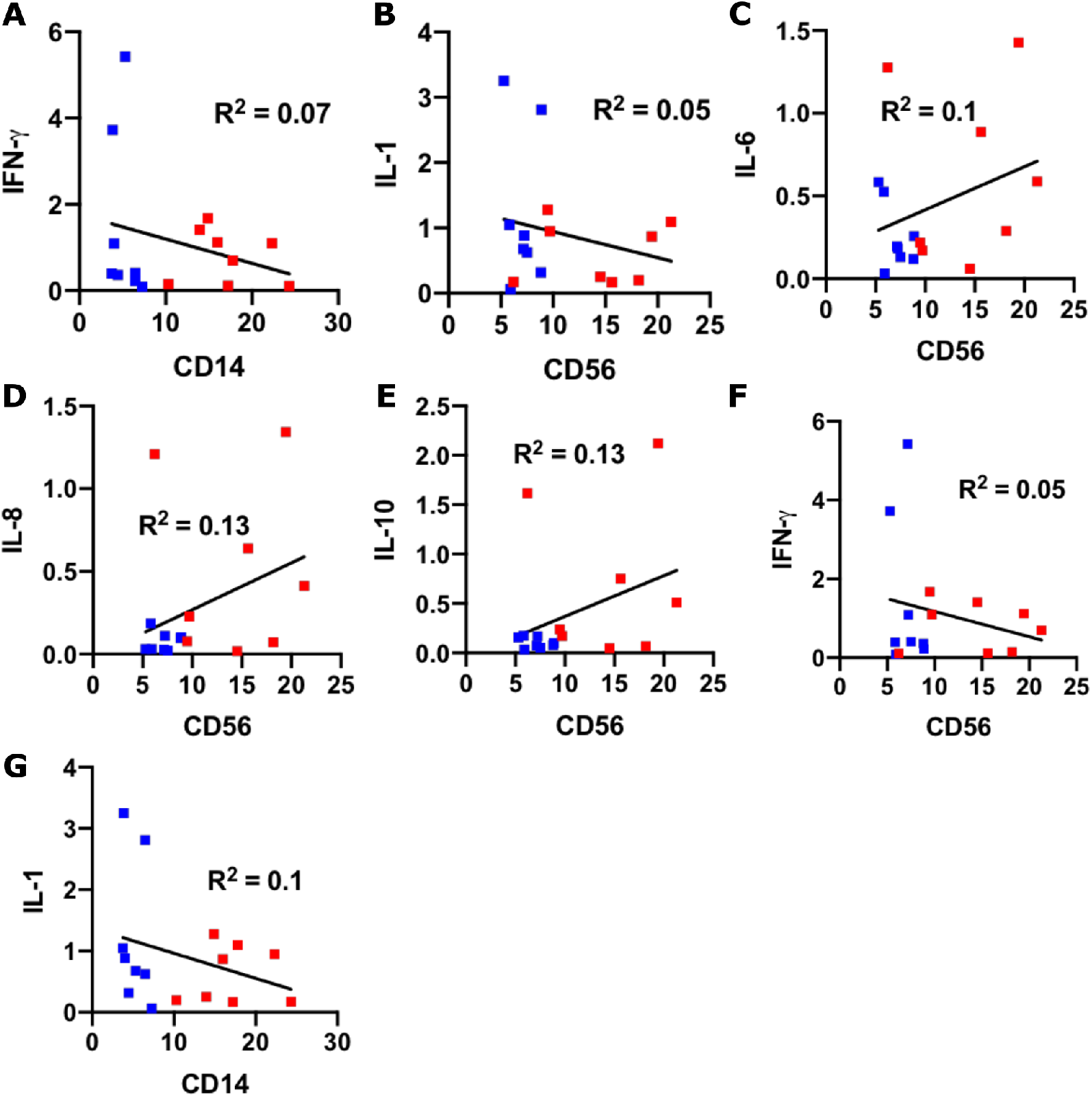
Non-significant correlations between cytokine abundance and immune cells. **(A-G)** Scatter plots of the indicated cytokines (Y axis; fold change relative to control) and immune cell markers (X axis; fluorescence ratio) for each placental sample (COVID-19+; n=16). Blue symbols = 2^nd^ trimester; red symbols = 3^rd^ trimester.

**Supplemental Figure 3:**
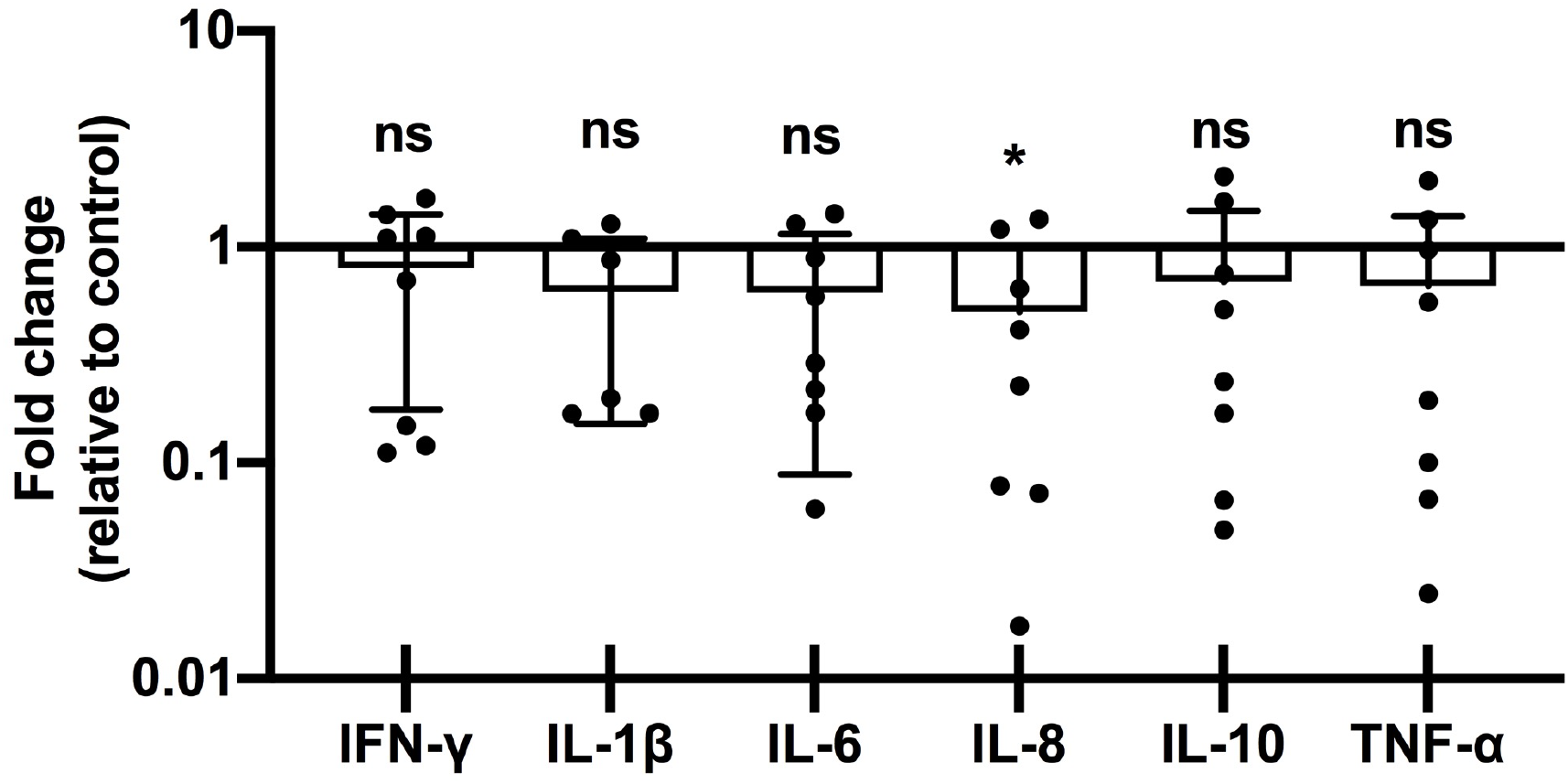
Placental cytokine expression in 3^rd^ Tri COVID group. Fold-changes in cytokine abundance for IFN-γ, IL-1β, IL-6, IL-8, IL-10, and TNF-α by qRT-PCR of placental decidua from women with 3^nd^ trimester COVID infection (n = 8) calculated as fold-change compared to control (n = 8). ns = not significant; * *p* <0.05.

